# ALS-linked KIF5A ΔExon27 mutant causes neuronal toxicity through gain of function

**DOI:** 10.1101/2022.03.05.483071

**Authors:** Devesh C. Pant, Janani Parameswaran, Lu Rao, Liang Shi, Ganesh Chilukuri, Zachary T. McEachin, Jonathan Glass, Gary J. Bassell, Arne Gennerich, Jie Jiang

**Author notes:** Correspondence should be addressed to Jie Jiang. These authors contributed equally to this work (Devesh C. Pant, Janani Parameswaran).

## Abstract

Mutations in the human kinesin family member 5A (*KIF5A*) gene were recently identified as a genetic cause of amyotrophic lateral sclerosis (ALS). Several KIF5A ALS variants cause exon 27 skipping and produce motor proteins with an altered C-terminal tail (referred to as ΔExon27). However, the underlying pathogenic mechanism is still unknown. In this study, we performed a comprehensive analysis of ΔExon27 at the single-molecule, cellular, and organism levels. Our results show that ΔExon27 is prone to form cytoplasmic aggregates and is neurotoxic. The mutation relieves motor autoinhibition and increases motor self-association, leading to drastically enhanced processivity on microtubules. Finally, ectopic expression of ΔExon27 in *Drosophila melanogaster* causes wing defects, motor impairment, paralysis and premature death. Our results suggest gain of function as an underlying disease mechanism in KIF5A-associated ALS.

## Introduction

Amyotrophic Lateral Sclerosis (ALS) is a progressive neurodegenerative disease characterized by loss of upper and lower motor neurons (MNs) leading to paralysis and death within 3–5 years after diagnosis (Chia *et al*, 2018). Most of ALS patients are classified as sporadic ALS (sALS), while about 10% of patients show a clear family history (fALS). Although varying in disease onset and duration, fALS and sALS patients cannot be differentiated by their clinical features. The mechanisms of disease pathogenesis leading to the exclusive demise of MNs remain unclear and there is no effective therapy. Over 30 genes have been linked to ALS, and several of these genes such as *PFN1* (profilin 1) and *TUBA4A* (tubulin, alpha 4A) are involved in cytoskeletal function and intracellular transport (Brenner & Weishaupt, 2019). Motor neurons are highly polarized cells with neurites that can exceed 1 meter in length. Various materials such as mRNA, proteins, lipids, membrane-bound vesicles, and organelles commute between cell bodies and synaptic terminals as they are transported along microtubules by two major families of motor proteins. Kinesin family (KIF) proteins mediate anterograde transport away from the cell body to neurite terminals whereas cytoplasmic dynein-1 drives the retrograde transport in the opposite direction. In 2018, two studies reported that mutations in KIF5A (OMIM# 602821) cause ALS in European and North American cohorts, further highlighting the role of intracellular transport disruption in disease pathogenesis (Brenner *et al*, 2018; Nicolas *et al*, 2018). Additional ALS patients with KIF5A mutations were subsequently identified in Asian cohorts (Faruq *et al*, 2019; Nakamura *et al*, 2021; Naruse *et al*, 2021; Zhang *et al*, 2019).

Human KIF5A, together with KIF5B and KIF5C, belongs to the kinesin-1 family and is one of the founding members of the kinesin superfamily with a total of 45 members (Hirokawa *et al*, 2009). The KIF5A motor complex is a heterotetramer consisting of two kinesin heavy chains (KHCs) and two kinesin light chains (KLCs). It uses the energy derived from ATP binding and hydrolysis to transport a variety of cargos processively (the ability to take hundreds of steps along microtubules before dissociation) to the plus-ends of microtubules. Studies in animal models support that KIF5A has essential functions in the development and functioning of the nervous system (Kanai *et al*, 2000; Tanaka *et al*, 1998; Xia *et al*, 2003). The KIF5A KHC contains an N-terminal catalytic motor domain, an α-helical stalk region, and a C-terminal tail region (**Fig. 1A**). KIF5A was previously identified as the causative gene for hereditary spastic paraplegia-10 (SPG10) and Charcot–Marie–Tooth type 2 (CMT2), with mutations mostly located in the N-terminal motor domain (Blair *et al*, 2006; Crimella *et al*, 2012; Fichera *et al*, 2004). Moreover, *de novo* heterozygous frameshift rare variants in the C-terminal domain of KIF5A were reported to be associated with neonatal intractable myoclonus (NEIMY), a severe infantile-onset neurologic disorder characterized by myoclonic seizures, hypotonia, dysphagia, and early development arrest (Duis *et al*, 2016; Rydzanicz *et al*, 2017). Interestingly, the ALS-associated variants reported in KIF5A are mainly clustered at the 5′ and 3′ splice junctions of exon 27 (Brenner *et al*., 2018; Nicolas *et al*., 2018). These variants cause a complete skipping of exon 27, yielding a protein with the normal C-terminal 34 amino acids replaced with 39 aberrant amino acids (referred to as ΔExon27 hereafter) (**Fig. 1A**).

**Fig. 1.**
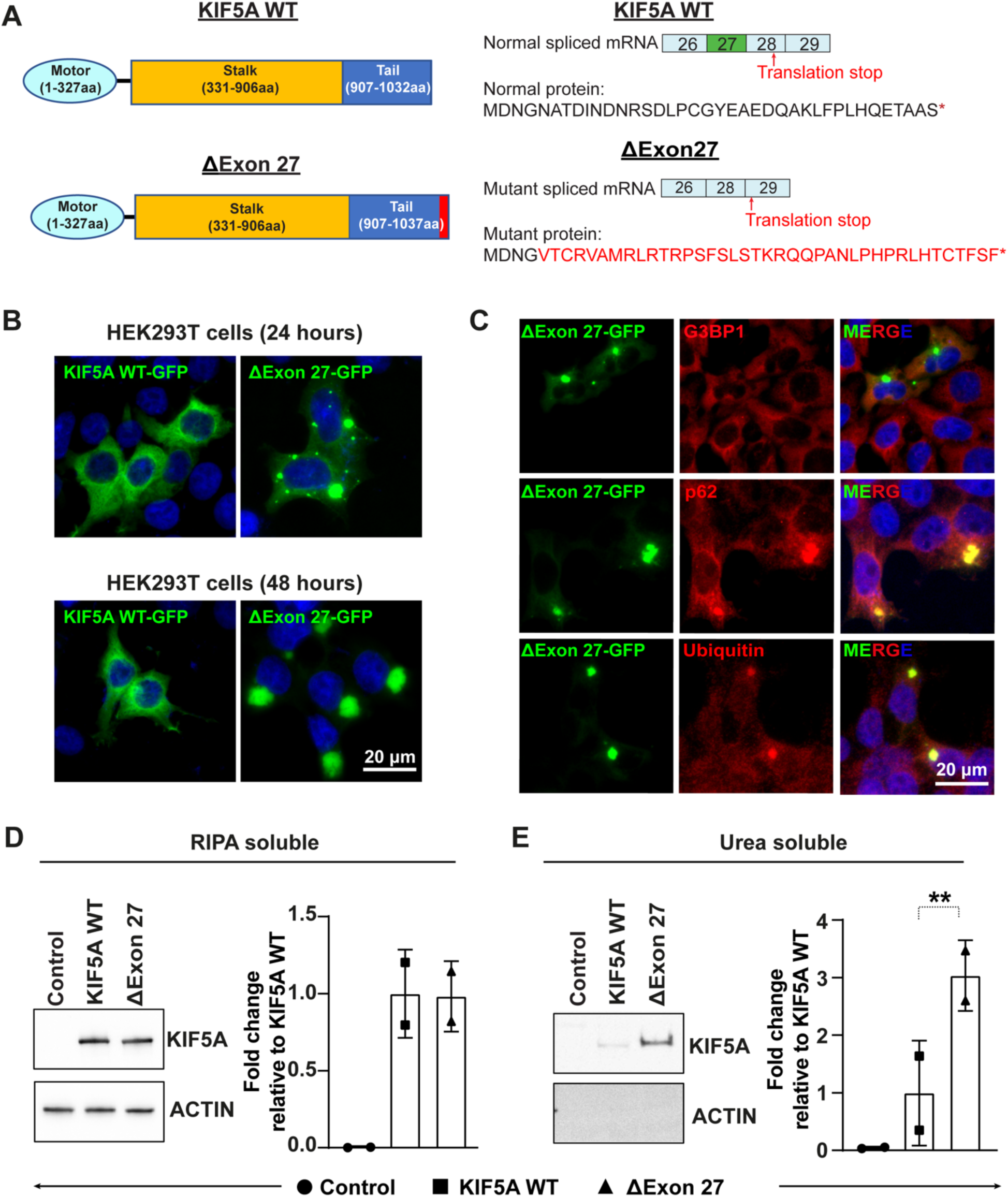
ALS-associated KIF5A ΔExon27 is prone to form cytoplasmic aggregates. **(A)** Schematic illustration of KIF5A WT and ΔExon27 showing the motor, stalk, and cargo domains (left). ALS-associated KIF5A variants disrupt the C-terminal tail by skipping exon 27 during mRNA splicing and produce a new C-terminal tail of 39 amino acids (red) replacing the normal tail of 34 amino acids (right). (**B**) HEK293T cells were transfected with KIF5A WT-GFP or ΔExon27-GFP for 24 and 48 hours. Nuclei were stained with DAPI. (**C**) Co-staining of G3BP1, p62, and Ubiquitin with KIF5A ΔExon27 granules. (**D-E**) Western blot analysis of RIPA soluble-fraction (D) and RIPA-insoluble, urea-soluble fraction of proteins prepared from HEK293T cells expressing empty plasmid (control), FLAG-KIF5A WT, FLAG-ΔExon27. Insoluble/soluble KIF5A fractions were detected with anti-FLAG and actin was used as a loading control. Data represent the mean ± SD.

Currently, the molecular mechanism of ΔExon27 underlying ALS pathogenesis is unclear. All known KIF5A ALS variants are autosomal dominant, and it is hypothesized that defective KIF5A alleles may lead to dysfunctional kinesin and impaired cargo transport. Alternatively, the altered C-terminal tail in the ΔExon27 mutant might confer detrimental effects through a toxic gain-of-function mechanism. In this study, we performed a comprehensive analysis of ΔExon27 using genetic, biochemical, and single-molecule methods. We find that ΔExon27 is prone to aggregation formation and is neurotoxic. Transgenic *Drosophila* expressing ΔExon27 display wing deficits, motor impairment, paralysis, and premature death. Mechanistically, ΔExon27 is constitutively active, displays increased motor self-association and drastically enhanced processivity on microtubules. Together, these results suggest the KIF5A ΔExon27 leads to neuronal toxicity via gain of function caused by constitutive activation and increased motor association and aggregation.

## Results

### KIF5A ΔExon27 is aggregation prone

We first expressed human wild type KIF5A (WT) and the ALS mutant ΔExon27 with C-terminal GFP tags in HEK293T cells. 24 hours after transfection, WT mainly diffuses in the cytoplasm (**Fig. 1B**), consistent with previous reports (Kamata *et al*, 2017; Yoo *et al*, 2019). Interestingly, almost 100% of cells expressing ΔExon27 show cytoplasmic granules of various sizes (**Fig. 1B**). At 48 hours, ∼4% WT expressing cells also showed a few granules, though significantly less abundant than those expressing ΔExon27, which usually have one single large inclusion in the cytoplasm (**Fig. 1B**). Similar results were also observed when these constructs were expressed in mouse neuroblastoma cells (N2a) (**Fig. S1A**). To exclude the possibility that this phenotype is due to the additional C-terminal GFP tag, we generated N-terminally FLAG-tagged KIF5A constructs. Like the C-terminally GFP-tagged ones, the N-terminally FLAG-tagged ΔExon27 also forms robust cytoplasmic granules in contrast to the homogeneously diffusive FLAG-tagged WT (**Fig. S1B**).

To identify the properties of these granules, we first determined whether these granules are positive for Ras GTPase-activating protein-binding protein 1 (G3BP1), a classical marker of stress granules that are implicated in ALS pathogenesis (Li *et al*, 2013). Few stress granules were observed in cells expressing either WT or ΔExon27 (**Fig. 1C, S1C**) and merely ∼2% ΔExon27 granules co-stain with G3BP1 (**Fig. S1D**), indicating that ΔExon27 granules are not stress granules. Protein aggregations are characteristic features of many neurodegenerative diseases. Because cytoplasmic inclusions of TAR DNA-binding protein 43 (TDP-43) were observed in >95% of ALS patients, (Neumann *et al*, 2006), we performed immunofluorescence staining to detect TDP-43. In cells expressing either WT or ΔExon27, TDP-43 remains within the nucleus (**Fig. S1E**). However, we found that ∼ 83% of ΔExon27 granules are positive for p62 and ubiquitin, common components of cytoplasmic inclusions found in many protein aggregation diseases (**Fig. 1C, S1D**). Immunoprecipitation of KIF5A using an antibody against the N-terminal FLAG tag also pulls down endogenous p62 only in cells expressing ΔExon27, but not WT (**Fig. S1F**). We further determined the solubility of WT and ΔExon27 in detergents of different strength. In the RIPA-soluble fraction, the expression level of ΔExon27 is comparable to that of WT, suggesting that the propensity of ΔExon27 to form cytoplasmic granules is not due to its higher expression (**Fig. 1D**). In the RIPA-insoluble and urea-soluble fraction, we detected significantly higher signal of ΔExon27 compared to WT (**Fig. 1E**). These results support that KIF5A ΔExon27 with the altered C-terminal tail is aggregation prone.

### ΔExon27 interacts with WT KIF5A and causes neuronal toxicity

KIF5A has been proposed to function as homodimers, with the dimerization domain located in the stalk region (Kanai *et al*., 2000). Since ALS-associated KIF5A variants are autosomal dominant, pathogenicity may be caused by mutant-WT KIF5A complexes. To mimic the patient scenario and determine whether ΔExon27 interacts with WT KIF5A, we co-expressed GFP-tagged WT and mApple-tagged ΔExon27 in HEK293T cells. WT and ΔExon27 colocalize near-perfectly, forming robust cytoplasmic aggregates 48 hours after transfection (**Fig. 2A**). In addition, immunoprecipitation of WT KIF5A pulls down ΔExon27 when the two constructs are co-expressed (**Fig. 2B**). We also expressed ΔExon27-GFP in N2a cells and stained for the endogenous KIF5A WT using an antibody generated with a peptide corresponding to the C-terminus of mouse KIF5A WT (amino acids 1007-1027), which does not recognize ΔExon27. While the endogenous KIF5A WT diffuses in the cytoplasm of non-transfected cells, it colocalizes with ΔExon27 aggregates in the transfected cells (**Fig. 2C**). These results suggest that KIF5A ΔExon27 interacts with WT and forms aggregates.

**Fig. 2.**
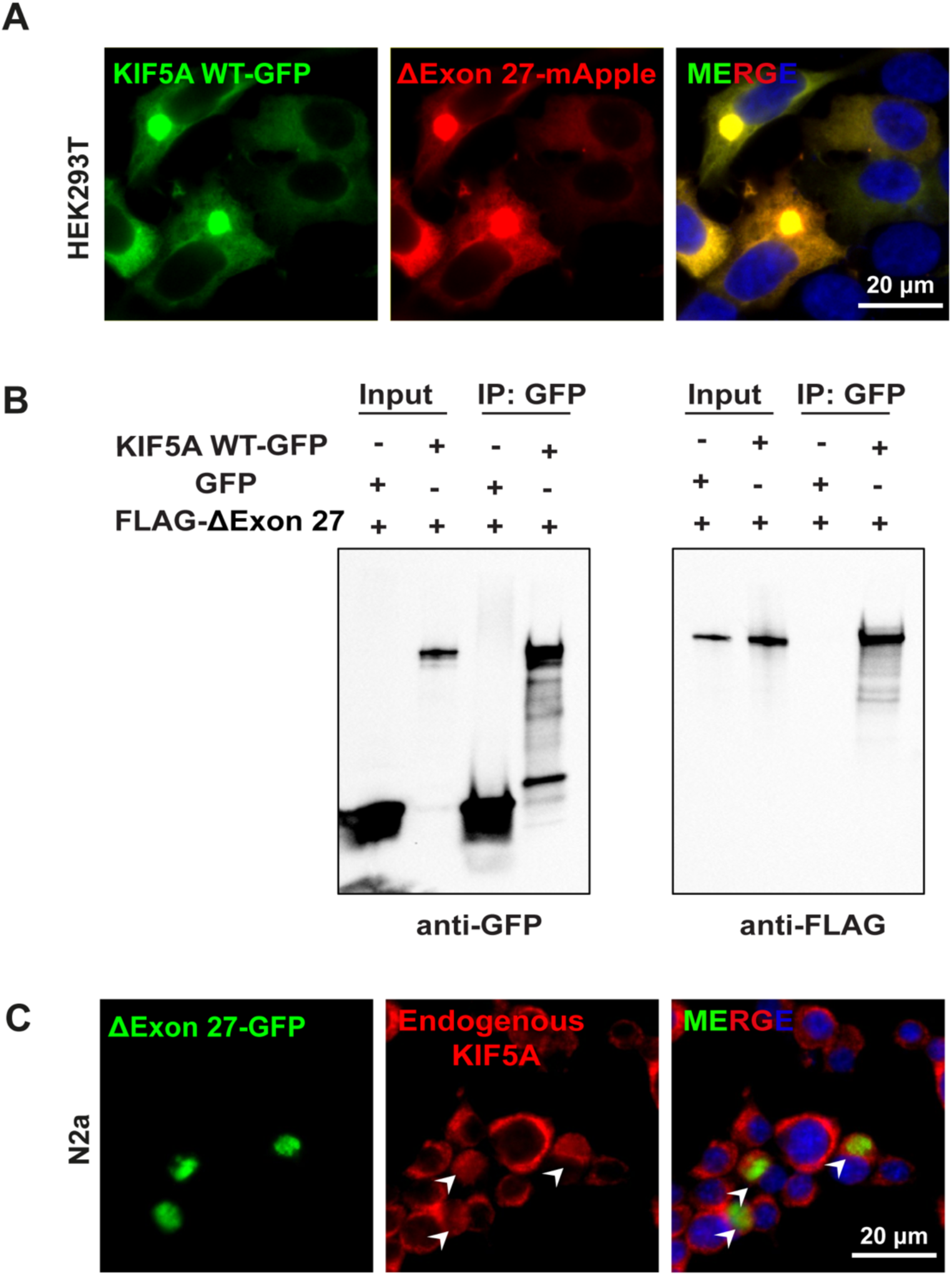
ΔExon27 interacts with WT KIF5A to form aggregates. **(A)** Expression of KIF5A WT and ΔExon27 tagged with GFP and mApple, respectively, in HEK293T cells. Images were taken 48 hours after transfection. **(B)** Pull down analysis showing the association of ΔExon27 with WT KIF5A. **(C)** Staining of mouse endogenous KIF5A in N2a cells transfected with ΔExon27-GFP using an antibody generated with a peptide corresponding to the C-terminus of KIF5A WT (amino acids 1007-1027), which does recognize ΔExon27. ΔExon27 granules are highlighted with arrowheads.

We further analyzed the expression pattern of WT and ΔExon27 in primary mouse cortical neurons. 24 hours after transfection, granules were observed in the soma of neurons expressing ΔExon27 and were even more abundant along neurites and at the terminal end. In contrast, WT KIF5A exhibits a diffuse pattern in the cytoplasm and along neurites (**Fig. 3A**). These granules are positive for p62, again suggesting they are likely caused by motor aggregations (**Fig. 3B**). We also co-stained these granules with the presynaptic marker, synapsin I, and observed neither colocalization nor close proximity of the granules to synapses (**Fig. S2A**). Given the robust accumulation of ΔExon27 along the neurites and at the terminal ends, we hypothesized that ΔExon27 preferentially accumulates at the plus-ends of microtubules. To test this, we obtained a highly polarized porcine kidney cell line (LLC-PK1) stably expressing tubulin-GFP (Rusan *et al*, 2001). Indeed, ΔExon27 granules were observed mainly at the peripheral extrusions (**Fig. S2B**) and colocalizes with microtubule plus-end-tracking protein EB1 (**Fig. S2C**).

**Fig. 3.**
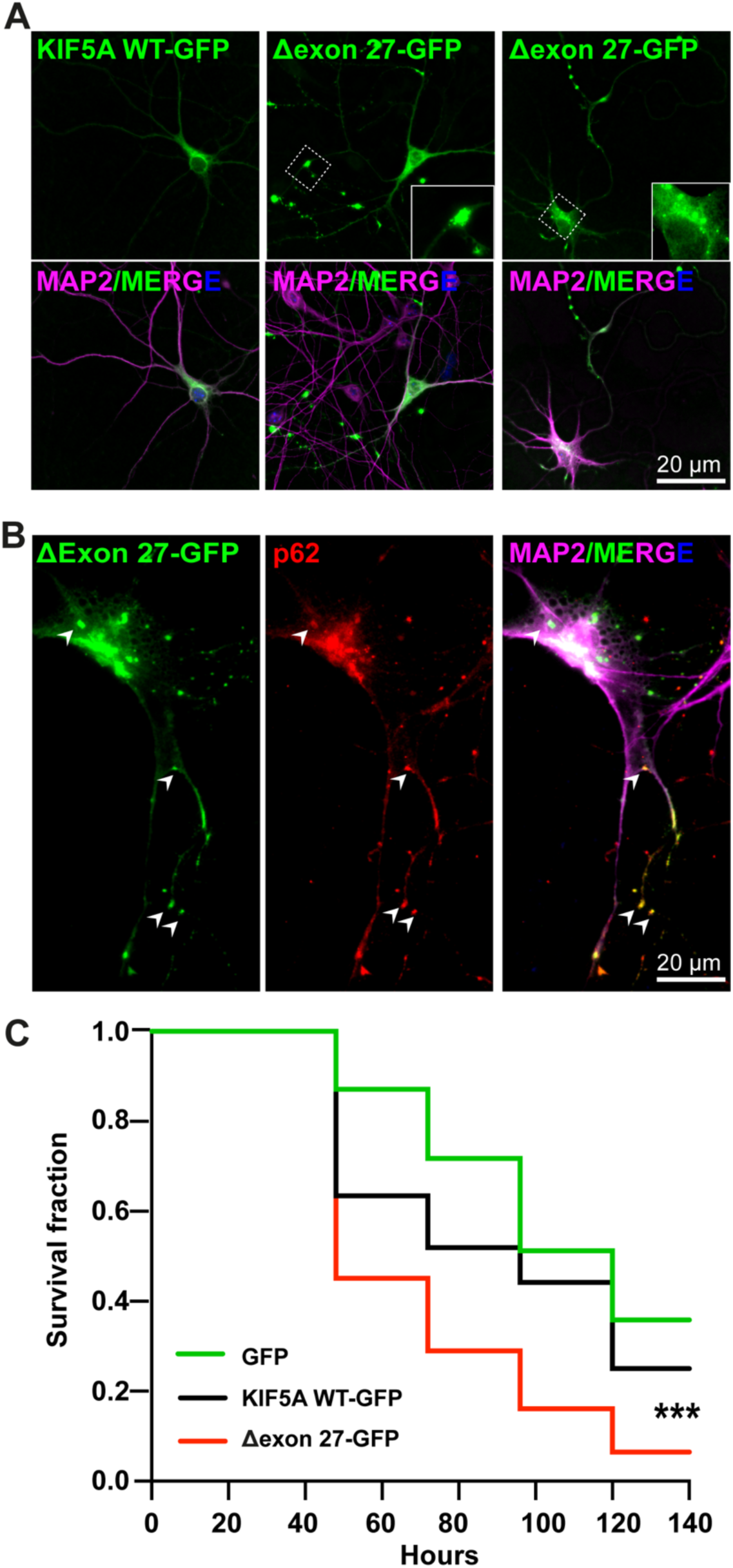
KIF5A ΔExon27 causes neuronal toxicity. (**A**) Mouse primary neurons transfected with KIF5A WT-GFP and ΔExon27-GFP were probed using anti-GFP antibody 24 hours after transfection. Neurons were identified by microtubule associated protein 2 (MAP2) and nuclei were stained with DAPI. **(B)** p62 staining in neurons expressing ΔExon27-GFP. Colocalization of p62 with ΔExon27 granules are highlighted with arrow heads. (**C**) 5-day-old mouse cortical neurons were transfected with mApple together with either GFP, KIF5A WT-GFP, or ΔExon27-GFP. Images were taken every 24 hours after transfection. Survival of neurons was analyzed by Kaplan-Meier survival analysis (*\*\*\*p* < 0.0001, log-rank test); 3 individual experiments were performed with similar results.

Protein inclusions formed by several disease-related proteins have been shown to trigger cellular toxicity (Soto & Pritzkow, 2018). To test whether aggregation-prone ΔExon27 causes neuronal toxicity, we co-transfected GFP, KIF5A WT-GFP, or ΔExon27-GFP together with mApple in mouse primary cortical neurons at DIV (days *in vitro*) 5 and used automated longitudinal microscopy to track the survival of hundreds of neurons indicated by mApple fluorescence over 6 days. The risk of death is significantly higher in neurons expressing ΔExon27-GFP than those expressing WT-GFP or GFP alone (**Fig. 3C**), indicating that ΔExon27 is neurotoxic.

### KIF5A ΔExon27 exhibits relieved autoinhibition, increased motor self-association, and enhanced processivity on microtubules

The accumulation of ΔExon27 granules at the plus-ends of microtubules suggests that the localization of granules observed in distal neurites may be driven by aberrant motility and/or association of the mutant protein along microtubules. Kinesin-1 motors transport cargoes by converting chemical energy from ATP to mechanical energy to drive processive movement on microtubules. When not transporting cargos, kinesin is autoinhibited to prevent ATP squandering. Autoinhibition is achieved by the C-terminal tail domain binding to the motor domain to keep it in a folded autoinhibited state (Dietrich *et al*, 2008; Kaan *et al*, 2011). Given that ΔExon27 possesses an altered C-terminal tail, we hypothesized that the autoinhibition of KIF5A might be compromised. To test this, we performed single-molecule total internal reflection fluorescence (smTIRF) microscopy assays to assess the mobility of three fluorophore-tagged KIF5A proteins: WT, a KIF5A construct in which the C-terminal cargo domain was removed rendering the motor incapable of autoinhibition (amino acids 1-906, referred to as ΔC, **Fig. S3A**), and ΔExon27. On microtubule filaments bound to the cover glass of the slide chamber, KIF5A WT motors mostly show non-motile interactions with the microtubule (**Fig. 4A**), suggesting the majority of motors are primarily in the autoinhibited state. The few WT motors that show motility display an average velocity of 0.45 ± 0.11 μm/s and processivity of 0.6 [0.57, 0.71] μm (median with 95 % CI). In both HEK293T and N2a cells, ΔC also shows diffused cytoplasmic expression like WT (**Fig. S3B**). As expected, and in contrast to WT, ΔC exhibits enhanced motility and significantly increased velocity (0.85 ± 0.07 μm/s) and processivity (1.63 [1.47, 1.81] μm) compared to WT as determined with the smTIRF assay (**Fig. 4C, Table 1**). Similar to ΔC, ΔExon27 motors also exhibits enhanced motility events along microtubules in the smTIRF assay, supporting the hypothesis that autoinhibition is relieved in ΔExon27. Intriguingly, while ΔExon27 displays increased velocity (0.66 ± 0.10 μm/s) compared to WT, its velocity is slower than ΔC and it exhibits drastically increased processivity (5.47 [3.84, 7.80] μm) relative to ΔC (*p* < 0.0001, **Fig. 4B, Table 1**). We also observed that the fluorescence intensities of moving ΔExon27 molecules were much stronger than those WT and ΔC (**Fig. S4**), indicating that ΔExon27 promotes self-association (or aggregation) among KIF5A motors and moves along the microtubules as multi-motor complexes. The recruitment of additional motors would increase the number of motor domains near the microtubules thereby decreasing the chance of complete dissociation of the motor complex from the microtubules during a run. Analysis of the initiation of motor movement indicates that ΔExon27 does not have a preference over the landing location on microtubules (**Fig. S3C**).

**Table 1.** Measured velocities and run lengths of all KIF5A constructs. The velocity data were fit with Gaussian distribution, and the run length data were fit with an exponential decay function. *Denotes the fluorophore with which KIF5A movement was tracked.

| Construct | Velocity (mean ± SD) ($\mu\text{m/s}$) | Processivity (lifetime [95% CI]) ($\mu\text{m}$) |
| --- | --- | --- |
| KIF5A WT-GFP | $0.45 \pm 0.11$ | 0.60 [0.51, 0.71] |
| $\Delta\text{Exon27}$-GFP | $0.66 \pm 0.10$ | 5.47 [3.84, 7.80] |
| $\Delta\text{C}$-GFP | $0.85 \pm 0.07$ | 1.63 [1.47, 1.81] |
| K998-GFP | $0.70 \pm 0.11$ | 2.07 [1.72, 2.50] |
| KIF5A WT-GFP/$\Delta\text{Exon27}$-mCherry (KIF5A WT-GFP*) | $0.62 \pm 0.10$ | 3.95 [2.81, 5.53] |
| KIF5A WT-GFP/$\Delta\text{Exon27}$-mCherry ($\Delta\text{Exon27}$-mCherry*) | $0.62 \pm 0.10$ | 4.49 [3.64, 5.63] |

**Fig. 4.**
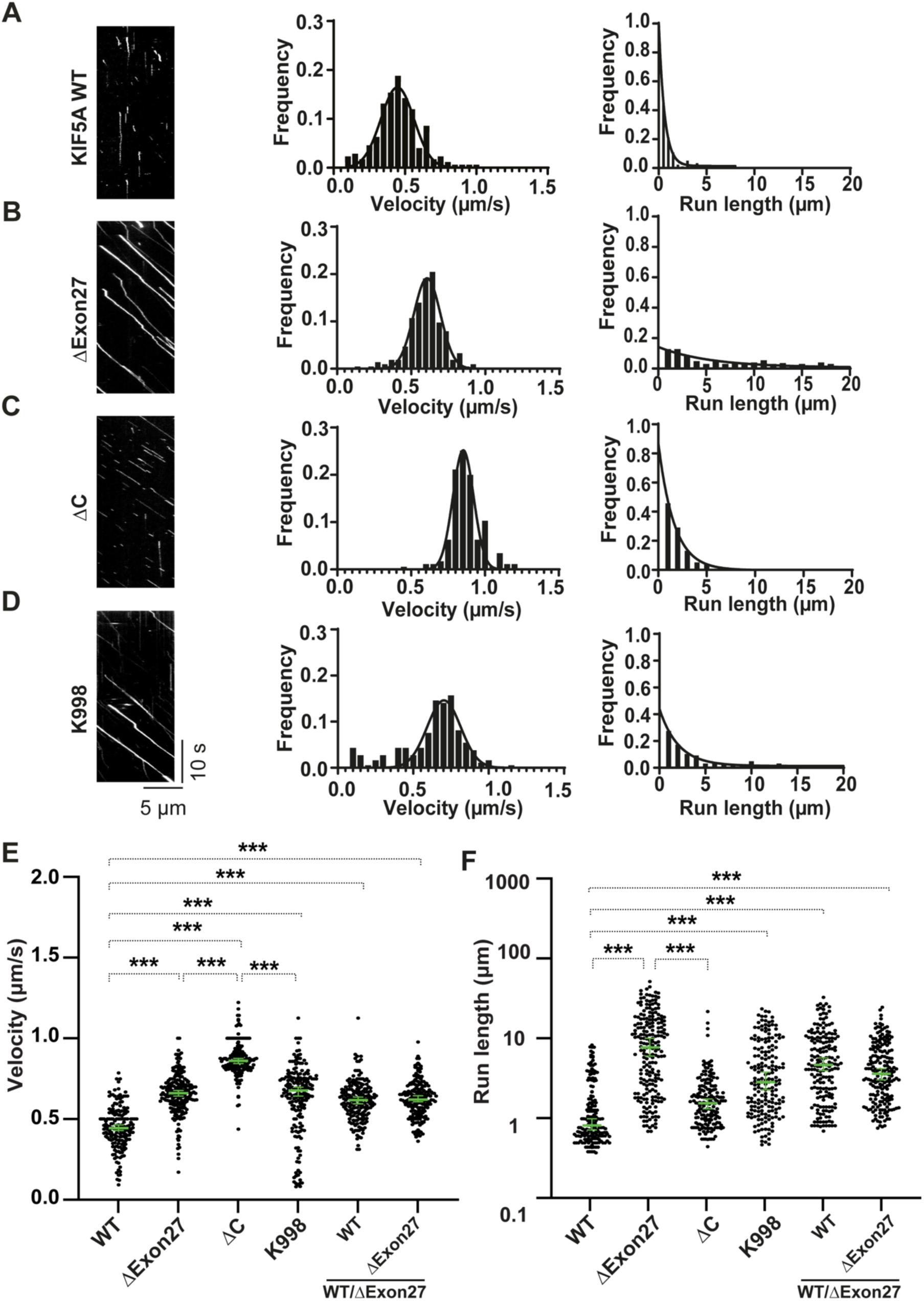
KIF5A ΔExon27 relieves motor autoinhibition and displays increased velocity and processivity. Kymographs and motility analyses of non-motile and mobile KIF5A proteins prepared from HEK293T cells. (**A**) KIF5A WT-GFP. From kymographs (example on the left), single-motor velocities (middle) and run lengths (right) were determined. (**B**) As in *A*, but for ΔExon27-GFP. (**C**) As in *A*, but for ΔC-GFP; (**D**) As in *A*, but for K998-GFP. The x-axis scale for the processivity graphs was limited to 20 μm to permit a direct comparison of the run lengths of the different constructs. Diagonal lines in the kymograph represent KIF5A molecules moving over time. The depicted scale bars are the same for all kymographs shown in this figure. The velocity data were fit with Gaussian distribution and the processivity data were fit with an exponential decay function. (**E-F**) Statistical analysis of the velocity **(E)** and processivity **(F)** of various KIF5A proteins. The green bars represent the median with 95% CI. The statistical comparison of velocity was performed using unpaired parametric t-test (*** *p* < 0.001). The run length is in log scale. The statistical evaluation of processivity was performed using ANOVA two way (\*\*\**p* < 0.001). The measured values for the velocities and run lengths are listed in Table 1.

To determine whether the new C-terminal tail of ΔExon27 is required for the drastically increased processivity, we made a construct that contains the same sequence region between WT and ΔExon27 (amino acids 1-998, referred to as K998, **Fig. S3A**). When expressed in HEK293T and N2a cells, K998 has diffused cytoplasmic localization as WT and ΔC (**Fig. S3B**). In the smTIRF assay, K998 displays a similar velocity (0.70 ± 0.11 μm/s) and processivity (2.07 [1.72, 2.50] μm) as those observed for ΔC (**Fig. 4D, a**nd **Table 1**). Importantly, the processivity of K998 is significantly lower than that of ΔExon27 (*p* < 0.0001), and the fluorescence intensity of motile motors is similar to that of WT and ΔC (**Fig. S4**). These results suggest that the new C-terminal tail confers a gain-of-function via the relief of KIF5A autoinhibition and promotion of motor self-association, which in turn causes KIF5A ΔExon27 motor complexes to move super-processively along microtubules.

Finally, we characterized the motility of WT and ΔExon27 complexes. WT and ΔExon27, being fluorescently tagged by GFP and mApple, respectively, were co-expressed in HEK293T cell. In the presence of ΔExon27, WT displays motile properties that resemble ΔExon27. Analysis using either WT-GFP or ΔExon27-mApple to assess motility yielded similar results for both velocity (0.62 ± 0.10 μm/s and 0.62 ± 0.10 μm/s, respectively) and processivity (3.95 [2.81, 5.53] μm and 4.49 [3.64, 5.63] μm, respectively) (**Fig. 4E-F, S3D**). Overlay of the kymograph of WT-GFP and ΔExon27-mApple demonstrates that most of the moving spots consist of both WT and ΔExon27 (**Fig. S3D**), consistent with our earlier observation that ΔExon27 and WT form complexes with each other (**Fig. 2**).

### Ectopic expression of KIF5A ΔExon27 in *Drosophila* leads to abnormal wings, motor deficits, paralysis, and premature death

To further investigate the role of KIF5A ALS mutant *in vivo*, we generated transgenic *Drosophila* expressing either WT or ΔExon27 with a C-terminal GFP tag. To avoid any influence of the genomic environment on transgene expressions, a single copy of *UAS>hKIF5A* transgenes were targeted to the well-established 68A4 landing site on chromosome III by PhiC31 integrase mediated insertion. These transgenes are not expressed at baseline but will be transcribed when Gal4 is introduced by genetic cross. The protein levels of WT and ΔExon27 are comparable when driven by the ubiquitously expressed tubulin-Gal4 (**Fig. S6A**). Constitutive ubiquitous expression of ΔExon27 in *Drosophila* causes immature-looking adult escapers with unexpanded wings in ∼40 % of flies and an early lethality (**Fig. 5A-C**). In contrast, neither the tubulin-Gal4 itself nor KIF5A WT causes any defects, confirming that the phenotype is exclusively due to ΔExon27 expression.

**Fig. 5.**
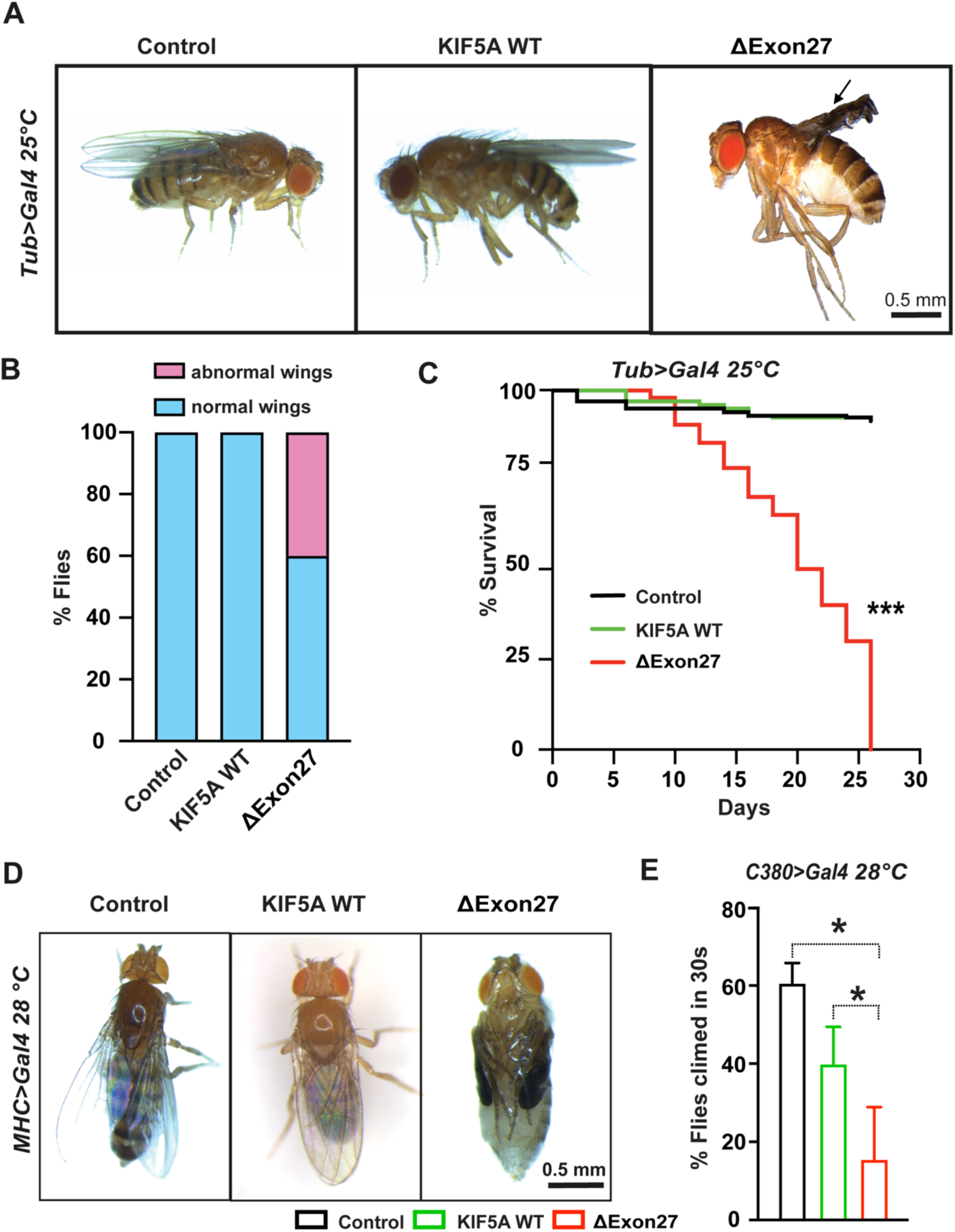
KIF5A ΔExon27 is toxic in *Drosophila melanogaster*. (**A**) Ubiquitous overexpression of ΔExon27 leads to wing defects (black arrows). (**B**) Percent of flies with normal and abnormal wings (n = 50 flies, 2 biological repeats). (**C**) Life spans were analyzed for flies expressing KIF5A WT-GFP and ΔExon27-GFP, and control flies expressing tubulin-Gal4 only. Survivorship was plotted over time. Expression of ΔExon27-GFP by the tubulin driver caused a substantial decrease in viability (*p* < 0.001, log-rank test). Survival curves represent an average of two trials. (**D**) Expression of ΔExon27-GFP in *Drosophila* muscles leads to complete paralysis. The pupal case of the fly expressing ΔExon27-GFP has been removed for this picture. The folded wings and legs are characteristic of the pupal state in flies expressing ΔExon27. (**E**) Negative geotaxis assay showing reduced motor function in 25-day-old flies expressing ΔExon27-GFP driven by a motor neuron specific driver (C380-Gal4) (n=50 each group). Bars indicate mean ± SD. Statistical analysis was performed using one way-ANOVA.

Since ALS is a neuromuscular degenerative disease, we confined the expression of transgenes in fly muscles using an MHC-Gal4 driver. Contrary to KIF5A WT or MHC-Gal4, all the ΔExon27-expressing flies fully develop to the pupal stage as pharate adults but are unable to hatch from their pupal cases when raised at 28 °C. After dissecting the ΔExon27-expressing flies from their pupal case, we found that they have no obvious defects other than what could be expected from being paralyzed with folded wings and legs, wet appearance, and no movement (**Fig. 5D**). When raised at 25 °C, ΔExon27-expressing flies can develop to adults and have similar expression levels as WT (**Fig. S6B**). However, they have dramatically reduced life span (**Fig. S6C**). We further expressed KIF5A transgenes specifically in motor neurons using the commonly used *Drosophila* motor neuron driver, C380-Gal4, and assessed their motor function with negative geotaxis assays. Flies expressing ΔExon27 show impaired climbing activity compared to those expressing KIF5A WT or control flies expressing C380-Gal4 **(Fig. 5E)**. Overall, our results highlight an imperative role of KIF5A ΔExon27 in mediating toxicity.

## Discussion

Two different types of KIF5A variants in ALS patients were initially highlighted: a single nucleotide variant (SNV) caused by a missense mutation (rs11347976 [c.2957C>T]) that changes amino acid 986 from proline to leucine, and variants predicted to affect splicing of exon 27 (Brenner *et al*., 2018; Nicolas *et al*., 2018). Whether the identified SNV represents a disease-causing or risk allele for ALS is unclear. Indeed, 11 of 29 patients carrying this SNV in the first study also carry genetic mutations in other known ALS genes (Brenner *et al*., 2018). In contrast, variants predicted to affect splicing of exon 27, although less common than rs11347976, are highly penetrant and were subsequently identified in multiple patient cohorts around the world (Faruq *et al*., 2019; Nakamura *et al*., 2021; Naruse *et al*., 2021; Zhang *et al*., 2019). These splicing variants were termed as “loss-of-function” mutations in the original studies since they led to loss of exon 27 at the RNA level and the replacement of the normal C-terminal 34 amino acids with a new tail of 39 amino acids at the protein level. However, the underlying disease mechanism of KIF5A ΔExon27 is currently unknown. Our results suggest that KIF5A ΔExon27 causes toxicity mainly through gain of function. ΔExon27 is particularly prone to form cytoplasmic aggregates compared to WT or KIF5A with the deletion of C-terminal tails. In addition, ΔExon27 is relieved from autoinhibition and displays drastically increased processivity through enhanced motor self-association caused by the altered C-terminal tail (**Fig. 6**). As a result, un-inhibited motors bind to microtubules and move processively toward the microtubule plus-ends and are therefore effectively removed from the pool of cytoplasmic motors needed for plus-end-directed cargo transport. It has been shown that homozygous *Kif5a* knockout in mice is embryonic lethal while heterozygous *Kif5a*^+/-^ animals are completely normal, indicating that having 50% of normal KIF5A levels is sufficient for its normal physiological functions (Xia *et al*., 2003). These results further support that KIF5A ΔExon27 causes ALS through a toxic gain-of-function mechanism.

**Fig. 6.**
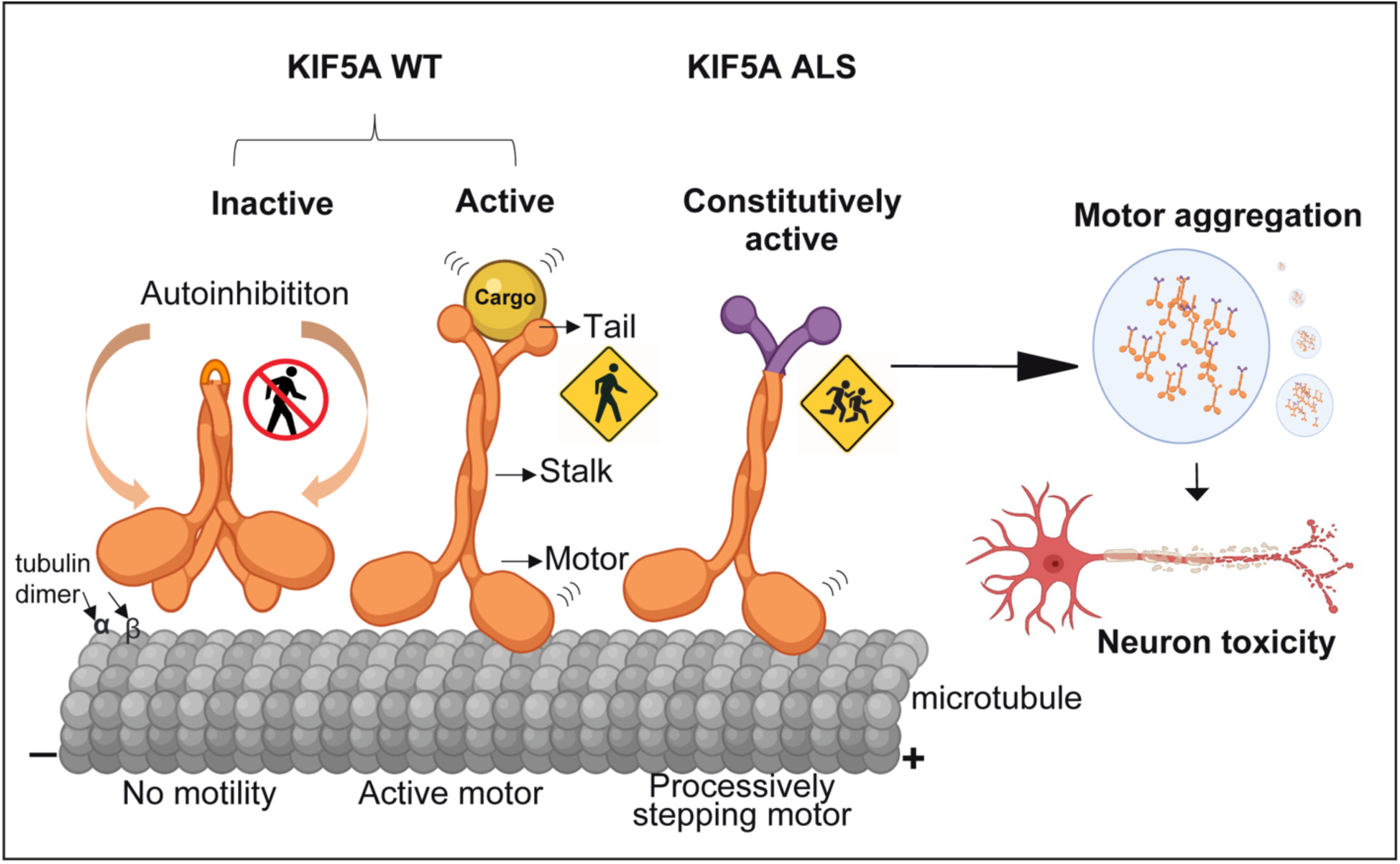
Proposed mechanisms of neuronal toxicity caused by KIF5A ΔExon27 gain of function. In the absence of tethered cargos, KIF5A WT is autoinhibited with the C-terminal tail binding to the N-terminal motor domain. When cargo binds to the C-terminal tail, the motor associates with microtubules and starts a processive run toward the plus-ends. ALS-associated KIF5A ΔExon27 with the new C-terminal tail relieves from the “autoinhibited” state even without cargo and self-associates to form multiple motors, leading to a drastically increased run-length on microtubules and accumulating at the plus-ends. ΔExon27 also forms complexes with WT KIFA and enhances motor self-association and aggregation.

In recent years, researchers have discovered that proteins bearing intrinsically disordered and low complexity domains coalesce into liquid droplets, a phenotype called liquid-liquid phase separation (LLPS). LLPS was initially recognized to play an important role in the formation and function of membraneless organelles such as Cajal bodies, stress granules and nucleolus, to name a few (Wang *et al*, 2021). It has also been shown to regulate local enrichment of molecules to activate cell signaling and acceleration of biochemical reactions (Sheu-Gruttadauria & MacRae, 2018; Su *et al*, 2016). By using seven different algorithms to predict proteins with intrinsic disordered domains, Seeger *et al*. showed that intrinsically disordered residues are present throughout the motor, stalk and tail domains of the kinesin family proteins. Strikingly, ∼71.8 % of kinesin tail domain residues are intrinsically disordered (Seeger & Rice, 2013). When expressed in mammalian cells, we found that KIF5A WT diffuses in the cytoplasm whereas ΔExon27 forms small granules at an earlier time point, which eventually increase in size likely by the fusion of multiple inclusions (**Fig. 1B**). It is interesting to note that similar but significantly fewer granules were also observed for KIF5A WT with time and continuous protein accumulation. Although it has been proposed that KIF5A functions as homodimers, whether KIF5A forms local condensates through LLPS and how this impact its role in transporting cargos warrants further study. Importantly, dysfunctional LLPS of DNA-and RNA-binding proteins such as TDP-43 and FUS have been implicated in ALS pathogenesis, leading to pathological protein aggregates in patients (Pakravan *et al*, 2021). We showed that ΔExon27 granules are positive for p62 and ubiquitin and are only soluble in stringent detergent such as urea (**Fig. 1C-E, Fig. S1C-D**). These data support that ΔExon27 is prone to form cytoplasmic aggregates that recruit p62-and ubiquitin-dependent proteolytic pathways.

Mutations of KIF5A have been previously linked to SPG10 and CMT2, two neurological diseases with neuronal axonopathy (Crimella *et al*., 2012). While all the ALS-related KIF5A variants are within the C-terminal tail, most SPG10/CMT2 mutations are located at the N-terminal motor region with a few in the stalk domain. The unique positions of KIF5A mutations that are associated with either SPG10/CMT2 or ALS provide an unprecedented opportunity to delineate pathogenic mechanisms underlying ALS. The kinesin-1 family of motor proteins are the best-understood members with respect to their mechanisms of motility at the single-molecular level (Gennerich & Vale, 2009). The fundamental regulatory mechanism for kinesin-1 is the transition from a “folded” (inhibited) to an “open” (activated) state (**Fig. 6**). In the folded state, the tail/cargo-binding domain binds to the motor domain (or “head”) and inhibits its activity by blocking the release of ADP (autoinhibition) (Hackney *et al*, 2009; Seeger & Rice, 2010). When the motor is activated presumably by cofactors or regulators, one of the two heads releases ADP upon microtubule binding, which starts a productive processive run (Gennerich & Vale, 2009; Hancock, 2016). Using single-molecule assays, we found that KIF5A ΔExon27 exhibits an increased velocity and a striking five-fold increase in processivity compared to KIF5A WT (**Fig. 4, Table 1**). Surprisingly, the processivity of ΔExon27 is much higher than ΔC, a construct where the C-terminal domain is deleted and autoinhibition is removed. This contrasts with the previous studies on the motile properties of SPG10 mutants, which exhibited reduced microtubule affinities and reduced velocities in microtubule-gliding assays (Ebbing *et al*, 2008). In addition, KIF5A WT and ΔExon27 interact to form oligomeric motors (**Fig. 2**) and display increased velocity and processivity like ΔExon27, suggesting that, when complexed, the mutant KIF5A confer its gain of function property to the WT protein. The consequence of this drastically increased processivity of KIF5A ΔExon27 to neuronal health is currently unknown. In primary cortical neurons, ΔExon27 is highly accumulated along the neurites and at the neurite terminals (**Fig. 3A-B**). Consistent with this, ΔExon27 granules are mainly observed at the peripheral extrusions of highly polarized LLC-PK1 cells and colocalize with microtubule plus-end-tracking protein EB1 (**Fig. S2B-C**). These studies suggest that enhanced processivity may result in an aberrant distribution of the motor along the neurites, with an increase in the local concentration of ΔExon27 near the microtubule plus-ends, which in turn could increase motor self-association and aggregation. In addition, kinesin proteins need to be transported back to the cell body by dynein/dynactin-based retrograde transport after delivering cargos to distal ends. It is thus possible that an imbalance between the movement of antegrade and retrograde motor proteins will eventually disrupt intracellular transport. Importantly, we demonstrated that K998 with the truncated tail behaves more like ΔC than ΔExon27. These results support the idea that the new C-terminal tail in ΔExon27 is required to confer gain of toxicity through the relief of autoinhibition and through self-association/aggregation of the mutant motors. In line with this notion, several single nucleotide deletion variants of KIF5A have also been identified in ALS patients (de Boer *et al*, 2021) which result in a C-terminal tail similar to that of the exon 27-skiping variants: Variants c2987delA (p.Asp996fs), c.2989delA (p.Asn997fs), and c.2996delA (p.Asn999fs) cause one nucleotide frame shifting at amino acid positions Asp996, Asn997, and Asn999 at the protein level, respectively (**Fig. S5**).

The identification of ALS-associated KIF5A mutations adds *KIF5A* to a growing list of known cytoskeletal-related genes implicated in ALS. We characterized ΔExon27-mediated toxicity in a transgenic *Drosophila* model. *Drosophila* is a versatile model system that has been widely used in the ALS field to study the molecular mechanisms of key biological functions given their genetic and overall experimental tractability (Liguori *et al*, 2021). We showed that ubiquitous expression of ΔExon27 in *Drosophila* cause wing abnormalities and shortened lifespan (**Fig. 5A-C)**. Tissue-specific expression revealed higher susceptibility to ΔExon27 expression in muscles and led to paralysis (**Fig. 5D**). Motor neuron specific expression of ΔExon27 also impairs fly climbing activities in negative geotaxis assays (**Fig. 5E**). The toxicity of ΔExon27 is further supported by observations that mouse cortical neurons expressing mutant KIF5A have reduced survival (**Fig. 3**). Additional genetic studies for disease modifiers of toxicity using our *Drosophila* KIF5A ALS models will likely yield new mechanistic insights of disease pathogenesis and identify therapeutic strategies for this detrimental disease.

## Materials and Methods

### DNA constructs

Human KIF5A cDNA was a gift from Dr. Gary Bassell (Emory University) and was used as a template to generate the following GFP (EmGFP), mApple and FLAG tagged constructs: KIF5A-GFP, KIF5A-mApple, FLAG-KIF5A [full length wild type (WT), ΔExon27, Δcargo (amino acids 1-906, ΔC), K998 (amino acids 1-998)]. Constructs used for *in vitro* studies were cloned into the pGW1-EmGFP backbone (a gift from Dr. Sami Bermada, University of Michigan) by PCR based cloning (Invitrogen). For longitudinal fluorescence microscopy pGW1-mApple was used. The human KIF5A WT and ΔExon27 were subcloned to the pUAST-attB backbone, a gift from Dr. Ken Moberg (Emory University), for the generation of transgenic *Drosophila*.

### *Drosophila* stocks and genetics

Fly crosses were maintained at 25°C in a humidified chamber with 12-hour light-dark cycles. Fly food was prepared with a standard recipe (water; cornmeal; yeast; agar; molasses; propionic acid). Parental stocks were maintained at room temperature. Transgenic flies *UAS>hKIF5A WT-GFP* and *UAS>hKIF5A ΔExon27-GFP* were generated by inserting the respective transgenes into the *attP2* site of *y*^*1*^ *w*^*67c23*^; *P[CaryP]attP*2 (BI #8622) by injection and *phiC31* integration (BestGene Inc, USA). The *tubulin-gal4* and *MHC-Gal4* line were generously provided by Dr. Ken Moberg (Emory University) and Dr. Udai Pandey (University of Pittsburgh), respectively. The motor neuron driver line *C380-Gal4* was purchased from Bloomington *Drosophila* Stock Center (BI # 80580). Flies were imaged using a Leica MC170 HD digital camera mounted on a Nikon SMZ800N stereo microscope.

### *Drosophila* survival assay

For the survival assay, parental flies were raised with standard food and under 12-hour day/night cycles at 25°C unless otherwise stated. The parents were allowed to mate and lay eggs for 3 days before discarded. The offspring from these parents were collected over a period of 24 hours and sorted by sex. Up to 12 male flies were kept in individual vials containing standard food. For each genotype, multiple replicate vials were set up so that the total sample size was 75 for each genotype. Flies were transferred onto fresh food every 2-3 days. The number of deaths was recorded each day. All survival assays were performed at 25°C.

### Negative geotaxis assay

Fly climbing activity was assessed using negative geotaxis assay as previously described (Nichols *et al*, 2012). Briefly, 15 flies were transferred, without anesthetization, into each plastic vial and placed in the apparatus. The vials were tapped down against the bench and the climbing was recorded on video for 1 minutes. The percent of flies climbing to the top of vials in 30s was determined manually in a blinded manner.

### Cell culture and transfection

Human embryonic kidney (HEK293T), and mouse neuroblastoma (N2a) cell lines from ATCC. The porcine kidney cell line (LLC-PK1) was a gift from Dr. Melissa Gardner (University of Minnesota). Cells were cultured in high glucose DMEM (Invitrogen) supplemented with 10% fetal bovine serum (Corning), 4 mM Glutamax (Invitrogen), penicillin (100 U/mL), streptomycin (100 μg/mL) and non-essential amino acids (1%). Cells were grown at 37°C in a humidified atmosphere with 5% CO_2_. Cells were transiently transfected using polyethylenimine (1 mg/mL) or Lipofectamine 2000 (Invitrogen). Experiments were performed either 24 or 48 hours after transfection.

### Primary cortical neuronal culture and transfection

Primary cortical neurons were prepared from C57BL/6J mouse embryos (Charles River) of either sex on embryonic day 17. Cerebral cortices were dissected and enzymatically dissociated using trypsin w/ EDTA (Thermo Fisher Scientific; 10 minutes), mechanically dissociated in Minimum Essential Media (MEM; Fisher) supplemented with 0.6% glucose (Sigma) and 10% Fetal Bovine Serum (FBS; Hyclone). Neurons were plated on coverslips (Matsunami Inc., 22 mm) or MatTek dishes coated with poly-l-lysine (Sigma). A total of 50,000 neurons were plated as a ‘spot’ on the center of the coverslip to create a small, high-density network. Neurons were cultured in standard growth medium [glial conditioned neurobasal plus medium (Fisher) supplemented with Glutamax (GIBCO) and B27 plus (Invitrogen)], and half of the media was exchanged 2-3 times a week until the experiment endpoints. No antibiotics or antimycotics were used. Cultures were maintained in an incubator regulated at 37 °C, 5% CO_2_ and 95% relative humidity as described (Valdez-Sinon *et al*, 2020). Cells were transiently transfected using Lipofectamine 2000 (Invitrogen) according to the manufacturer’s instructions.

### Longitudinal fluorescence microscopy

Mouse primary cortical neurons were transfected with mApple and various KIF5A constructs and imaged by fluorescence microscopy at 24 hour intervals for 7 days as described (Weskamp *et al*, 2019). Custom scripts were used to automatically generate region of interest corresponding to each cell and determine time of death based on rounding of the soma, retraction of neurites, or loss of fluorescence. The time of death for individual neurons was used to calculate the risk of death in each population relative to a reference group. Images were acquired using Keyence BZ-X810 microscope with a 10× objective and analyzed by Image J. The images were stitched and stacked, and cell death was scored using the criteria mentioned above.

### Immunofluorescence

Cells were fixed in 4% paraformaldehyde (Electron Microscopy Sciences) for 20 minutes, washed three times for 5 minutes with phosphate buffer saline (1× PBS, Corning) and treated with 0.1% Triton ×-100 (Sigma) in PBS. Cells were blocked for 30 minutes in blocking solution consisting of 4% bovine serum albumin (Sigma) in PBS. Cells were incubated overnight in primary antibodies: rabbit anti-G3BP1 (Proteintech), rabbit anti-p62 (Proteintech), mouse anti-p62 (Novus Biologicals), rabbit anti-TDP-43 (Proteintech), rabbit anti-synapsin I (Sigma), guinea pig anti-MAP2 (Synaptic Systems) diluted in blocking solution (1:500). The next day, cells were washed 3 times for 5 minutes in PBS and incubated in secondary antibodies in blocking solution for one hour at room temperature. After washing 3 times for 5 minutes, coverslips with the cells were mounted using Prolong Gold Antifade mounting media (Invitrogen). Images were acquired with Keyence BZ-X810 microscope with a 60× oil objective. For image analysis, around 100-150 transfected cells were counted for each genotype in each experiment. Quantification of colocalization of ΔExon27 aggregates with G3BP1, p62, and ubiquitin were performed manually in a blinded manner using ImageJ analysis.

### Western Blotting and Immunoprecipitation

Whole cell extracts were isolated using RIPA Lysis Buffer pH 7.4 (Bio-world, USA) supplemented with Halt™ protease and phosphatase inhibitor cocktail (ThermoFisher Scientific), followed by DNA shearing. After centrifuge, the pellet was washed for 3 times and dissolved in 8 M urea buffer (10 mM Tris, pH 8.0; 8 M urea). The concentration of the isolated proteins was determined using BCA Protein Assay Reagent (Pierce, USA). 20 μg of protein was resolved in 4-20% precast polyacrylamide gel (Bio-Rad, USA). For immunoprecipitation, HEK 293T cells were collected and lysed using Pierce™ IP Lysis Buffer (ThermoFisher Scientific) supplemented with Halt™ protease and phosphatase inhibitor cocktail (ThermoFisher Scientific), and 500 μg of protein lysate was precleared with 25 μl of Chromotek GFP-Trap® or Pierce™ Anti-FLAG magnetic agarose beads. Following the manufacturer’s instructions, protein lysate was immunoprecipitated for 2 h at 4°C with the following antibodies: mouse anti-GFP (1:1000; Takara Bio), mouse anti-FLAG (1:1000; Sigma). The input 10% was analyzed with Western blot. Proteins were transferred to nitrocellulose membranes (0.2 μm, Bio-Rad) and incubated with primary antibodies: mouse anti-GFP (1:2000; Clontech), mouse anti-FLAG (1:1000; Sigma), rabbit anti-KIF5A N-terminus (1:1000; GeneTex), rabbit anti-KIF5A C-terminus (1:1000; abcam), rabbit anti-β actin (1:2000; GeneTex) overnight at 4°C followed by HRP-conjugated secondary antibodies (ABclonal) or IRDye secondary antibodies (Li-cor) at room temperature for 1 hour. Super Signal West Pico (Pierce, USA) was used for detection of peroxidase activity. Molecular masses were determined by comparison to protein standards (Thermo Scientific). Band intensities were measured using ImageJ and normalized to tubulin or actin.

### Generation, expression, and purification of K490-EmGFP

K490-EmGFP gene fragment was generated by PCR with 25 bp overlap with a modified NEB SNAP-tag backbone in which the SNAP-tag was replaced by a C-term 6His tag. The plasmid was generated by Gibson assembly. The insertion was confirmed by enzymatic digestion and agarose gel electrophoresis. Protein expression and purification were performed as previously described (Budaitis *et al*, 2021). Briefly, the plasmid was transformed into BL21-CodonPlus (DE3)-RIPL competent cells (Agilent, #230280). A single colony was inoculated in 1-mL TB medium (Laboratory, 2006) with 50 μg/mL chloramphenicol and 25 μg/mL carbenicillin. After shaken at 37 °C overnight, the 1 mL culture was inoculated into 400 mL TB medium and was shaken at 37 °C for 5 hours. The cell culture was then cool down on ice to <18°C, and IPTG was added to final 0.1 mM concentration. The protein expression was induced at 18°C overnight with vigorous shaking. Afterwards, the cell was harvested by centrifugation at 3000 *g* for 10 min. The supernatant was discarded, and the pellet was resuspended in 5 mL B-PER™ complete bacterial protein extraction reagent (Thermo Scientific, #89821) supplemented with 2 mM DTT, 0.2 mM ATP, 4 mM MgCl_2_, 2 mM EGTA, and 2 mM PMSF. The resuspended solution was flash frozen and stored at –80 °C before purification.

To purify K490-EmGFP, the cell solution was thawed at 37 °C, followed by nutation at room temperature for 20 min. The cells were further lysed *via* douncing for 10 strokes on ice, and then cleared by centrifuging at 80000 rpm for 10 min at 4 °C using a Beckman Coulter tabletop centrifuge. At the same time, 0.5 mL Roche Ni-NTA resin (MilliporeSigma, #5893682001) was washed with 2×1 mL of wash buffer [50 mM HEPES, 300 mM KCl, 2 mM MgCl_2_, 1 mM EGTA, 10 % glycerol, 1 mM DTT, 0.1 mM ATP, 1 mM PMSF, 0.1 % Pluronic F-127 (w/v), pH 7.2]. The cleared lysate was carefully added to the Ni-NTA resin and allowed to flow through. The resin was then washed with 5×2 mL wash buffer. The protein was eluted with elution buffer [50 mM HEPES (pH 7.2), 150 mM KCl, 2 mM MgCl_2_, 1 mM EGTA, 250 mM imidazole (pH 8.0), 1 mM DTT, 0.1 mM ATP, 1 mM PMSF, 0.1% Pluronic F-127 (w/v), 10% glycerol].

### Microtubule polymerization

Microtubule polymerization was performed as described (Rao *et al*, 2018). Briefly, 2 μL of 1 mg/mL Cy5-labeled tubulin (prepared as described in (Nicholas *et al*, 2014)), 2 μL of 1 mg/mL biotinylated tubulin (Cytoskeleton Inc., #T333P), and 2 μL of 10 mg/mL unlabeled tubulin (Cytoskeleton Inc., #T240) was mixed on ice. 0.6 μL of 10 mM GTP was added to the tubulin solution, and the mixture was incubated at 37 °C for 15 minutes. Then 0.72 μL of 0.1 mM taxol in DMSO was added to the mixture, which was further incubated at 37 °C for 15 minutes. Tubulin that was not incorporated into the microtubules were removed by centrifugation through a glycerol cushion [BRB80 (80 mM PIPES, 2 mM MgCl_2_, 1 mM EGTA) with 60% (v/v) glycerol, 10 μM taxol, and 1 mM DTT, pH 6.8]. The pellet was re-suspended in resuspension buffer [BRB80 with 10% (v/v) glycerol, 10 μM taxol, and 1 mM DTT, pH 6.8] to final concentration of 1 mg/mL microtubules. The microtubule solution was stored at room temperature in the dark for several days.

### Single-molecule total internal reflection fluorescence (smTIRF) microscopy assay

HEK293T cells were grown in 100 mm tissue culture dish (VWR). After 48 hours post transfection, cells were washed with sterile PBS and detached from the dish using a cell scraper. Cells were resuspended in ice cold RIPA lysis buffer supplemented with Halt™ protease and phosphatase inhibitor cocktail (Thermo Fisher Scientific). Lysates were centrifuged at 17,000 *g* for 5 minutes at 4 °C. The supernatant was aliquoted, flashed frozen in liquid nitrogen, and stored at – 80 °C. smTIRF assay was carried out similarly as described before (Rao *et al*., 2018). Briefly, a coverslip (No. 1.5, Zeiss, #474030-9000-000) was cleaned by ethanol, and a flow chamber was assembled using the cleaned coverslip, a glass slide, and two stripes of cut parafilm as described before (Nicholas *et al*., 2014). 0.5 mg/mL of BSA-biotin was flown into the chamber, and the slide was incubated in a humidity box for 5 minutes. Afterwards, the chamber was washed with 3 × 20 μL of blocking buffer (BB) [BRB80 with 1% Pluronic F-127 (w/v), 10 μM taxol, 2 mg/mL BSA, 1 mg/mL α-casein, pH 6.8], and incubated for 10 minutes in the humidity box. After the surface was passivated, 0.25 mg/mL of streptavidin was flown into the chamber and incubated for 5 minutes. The chamber was then washed with 3 × 20 μL BB. 0.5 μL of 0.2 mg/mL microtubules was diluted in 19.5 μL BB and subsequently flown through the chamber. The chamber was washed with 2 × 20 μL BB and 20 μL motility buffer (MB) [60 mM HEPES, 50 mM KAc, mM MgCl_2_, 1mM EGTA, 0.5% Pluronic F-127 (w/v), 10% glycerol (v/v), 10 μM taxol, 1 mM DTT, 2 mg/mL BSA, 1 mg/mL α-casein, pH 7.2]. The final assay solution was prepared by mixing 3 μL of 50 mg/mL BSA, 1 μL of 100 mM ATP, 2 μL of 25× protease inhibitor cocktail (Roche, #4693159001), 1 μL of 50 mM biotin, 1 μL of appropriated diluted HEK293T cell lysate or K490, and 42 μL of MB. The final dilution of the lysate was 200× to 1000× depending on the construct. The final dilution of K490 was 10000× (200 pM). 2 × 20 μL of the assay solution was flown through the chamber, and the chamber was sealed by vacuum grease. The images were acquired by 200 ms/frame. The data was analyzed using a home-built MATLAB program.

### Statistical analysis

Statistical analyses and graphs were prepared in GraphPad Prism (version 9). Data are expressed as mean ± SD or mean± SEM as shown in figure legends. Student t-test or one-way ANOVA was used for statistical analysis unless specified in figure legends. *p* less than 0.05 was considered significant (\**p* < 0.05, \*\**p* < 0.01, \*\*\**p* < 0.001).

## Acknowledgements

We thank members of the Jiang and Bassell labs for their helpful discussions. We also thank lab members of Drs. Kenneth Moberg and Victor Faundez for their technical help and discussions.

## Author contributions

DCP, JP, and JJ performed and analyzed all *in vivo* and cell-based experiments. LR performed and analyzed the single-molecule motility assays. AG designed and interpreted single-molecule data and edited the manuscript. GC helped in data analysis and *Drosophila* experiments. ZTM and JG provided biological reagents. DCP, JP, and JJ wrote the manuscript.

## Data and materials availability

All data are available in the main text or the supplementary materials.

## Supplementary Figures

## Supplementary Materials for

**Fig. S1.**
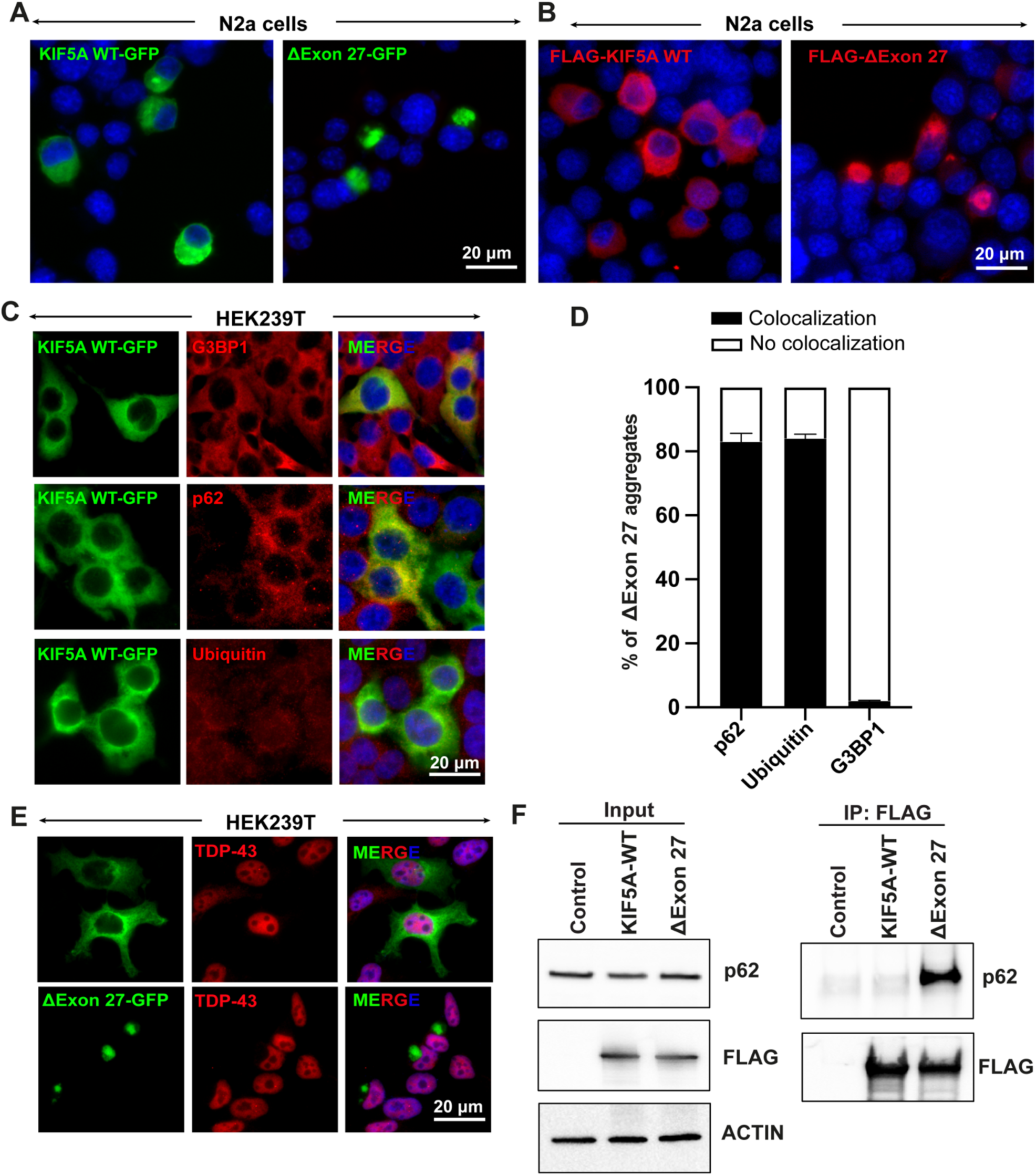
KIF5A ΔExon27 is prone to form cytoplasmic aggregates. **(A-B)** Expression of human KIF5A WT and ΔExon27 either with C-terminal GFP **(A)** or N-terminal FLAG tags **(B)** in N2a cells. **(C)** Co-localization of G3BP1, p62, and Ubiquitin markers with KIF5A WT expressed in HEK293T cells. (**D**) Percent of cells with G3BP1, p62 or Ubiquitin markers colocalized with cytoplasmic Δexon27 granules. **(E)** Staining of TDP-43 in HEK293T cells expressing either KIF5A WT or Δexon27 showed nuclear TDP-43. (**F**) Immunoprecipitation of KIF5A using antibodies against the N-terminal FLAG tag also pulled down endogenous p62 only in cells expressing Δexon27, but not WT KIF5A.

**Fig. S2.**
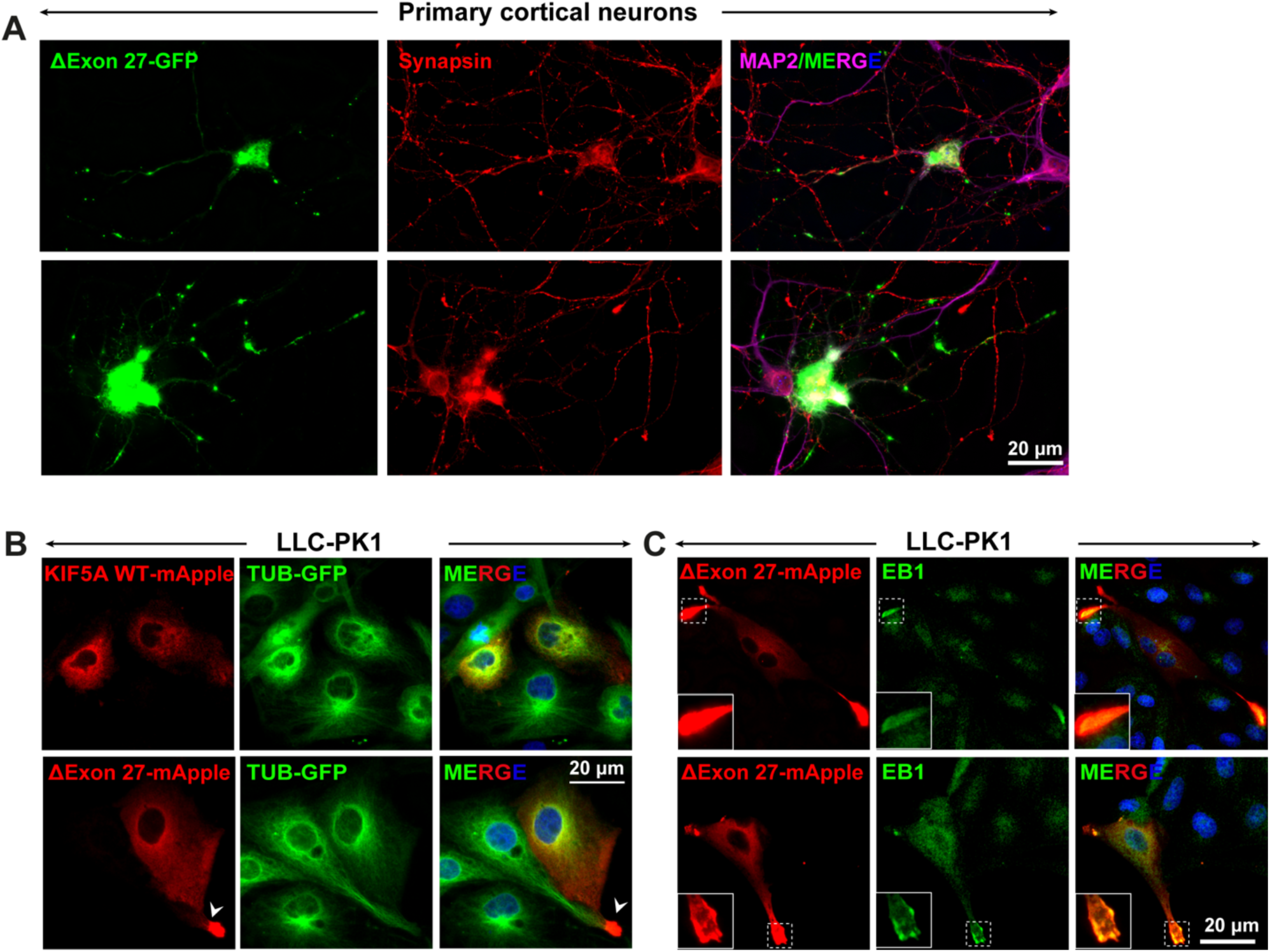
KIF5A ΔExon27 accumulates at the plus-ends of microtubules. **(A)** Primary neurons expressing KIF5A Δexon27 stained with pre-synaptic marker synapsin I. (**B**) Expression of KIF5A WT-mApple and Δexon27-mApple in LLC-PK1 cells stably expressing tubulin-GFP. KIF5A Δexon27 showed aggregation at proximal tubule regions as shown by white arrowhead. (**C**) Δexon27 expressed in LLC-PK1 cells showed colocalization with microtubule plus-end-tracking protein EB1 at the proximal extrusions.

**Fig. S3.**
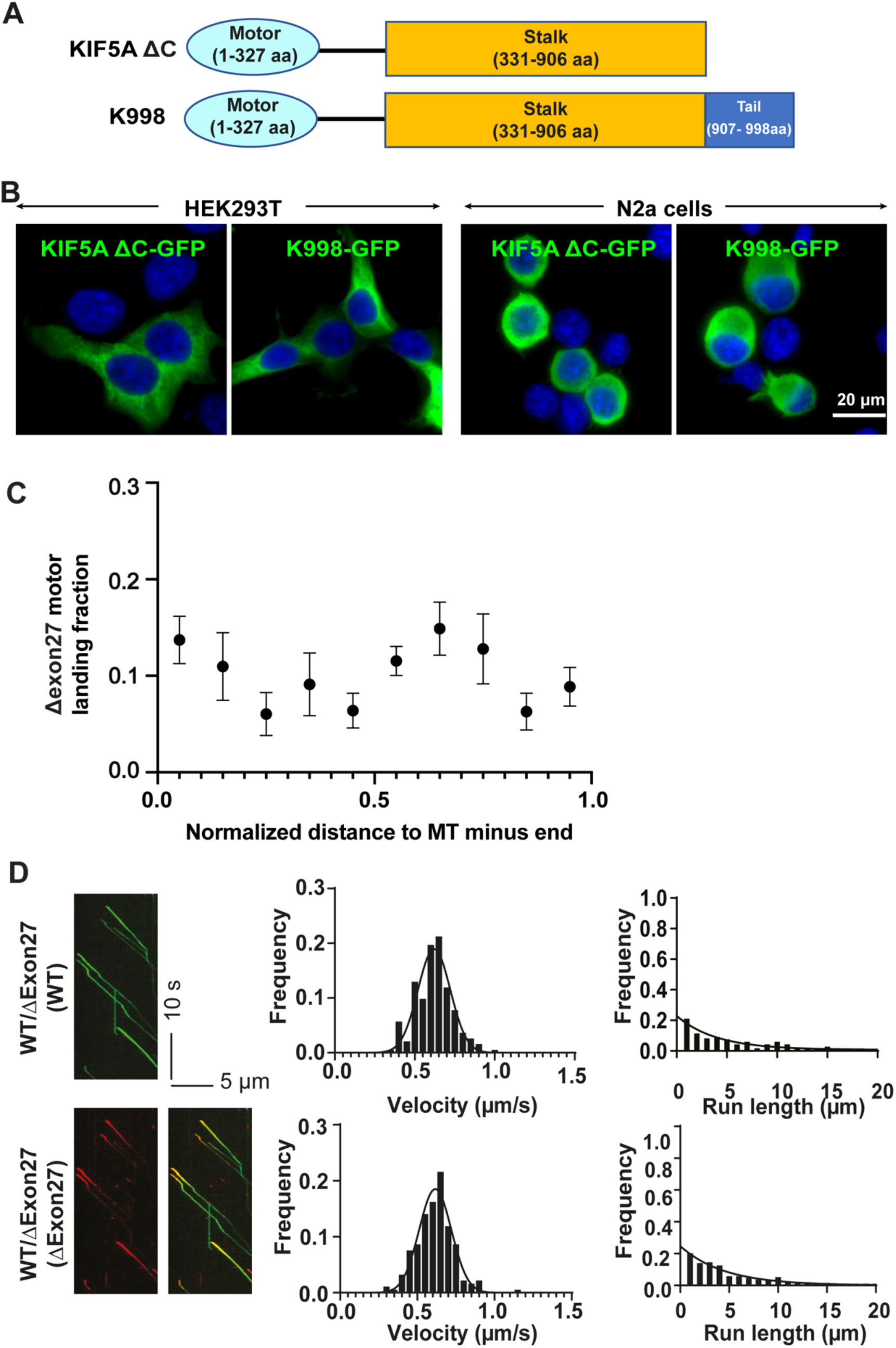
Motile properties of KIF5A ΔExon27 and characterization of truncation mutants K998 and ΔC. **(A)** Schematic illustrations of KIF5A truncated mutants, K998 and ΔC. **(B)** Both KIF5A K998 and ΔC truncated proteins diffuse in the cytoplasm when expressed in HEK293T and N2a cells. **(C)** Motor landing rate of KIF5A ΔExon27 along the microtube was accessed. ΔExon27 does not have a preference over the landing location on microtubules. **(D)** Motor velocities (middle) and run lengths (right) were determined based on kymographs (example on the left) for KIF5A WT and ΔExon27 complexes. HEK293T cells were transfected with both KIF5A WT-GFP and ΔExon27-mApple. The movements of WT and ΔExon27 were assessed by tracking either GFP or mApple, respectively. The right kymograph in *D* is the overlay of the GFP and mApple of moving WT-GFP/ΔExon27-mApple complexes. The x-axis scale for the processivity graphs was limited to 20 μm to permit a direct comparison of the run lengths of the different constructs. Diagonal lines in the kymograph represent KIF5A molecules moving over time. The velocity data were fit with Gaussian distribution and the processivity data were fit with an exponential decay function. The measured values for the velocities and run lengths are listed in Table 1.

**Fig. S4.**
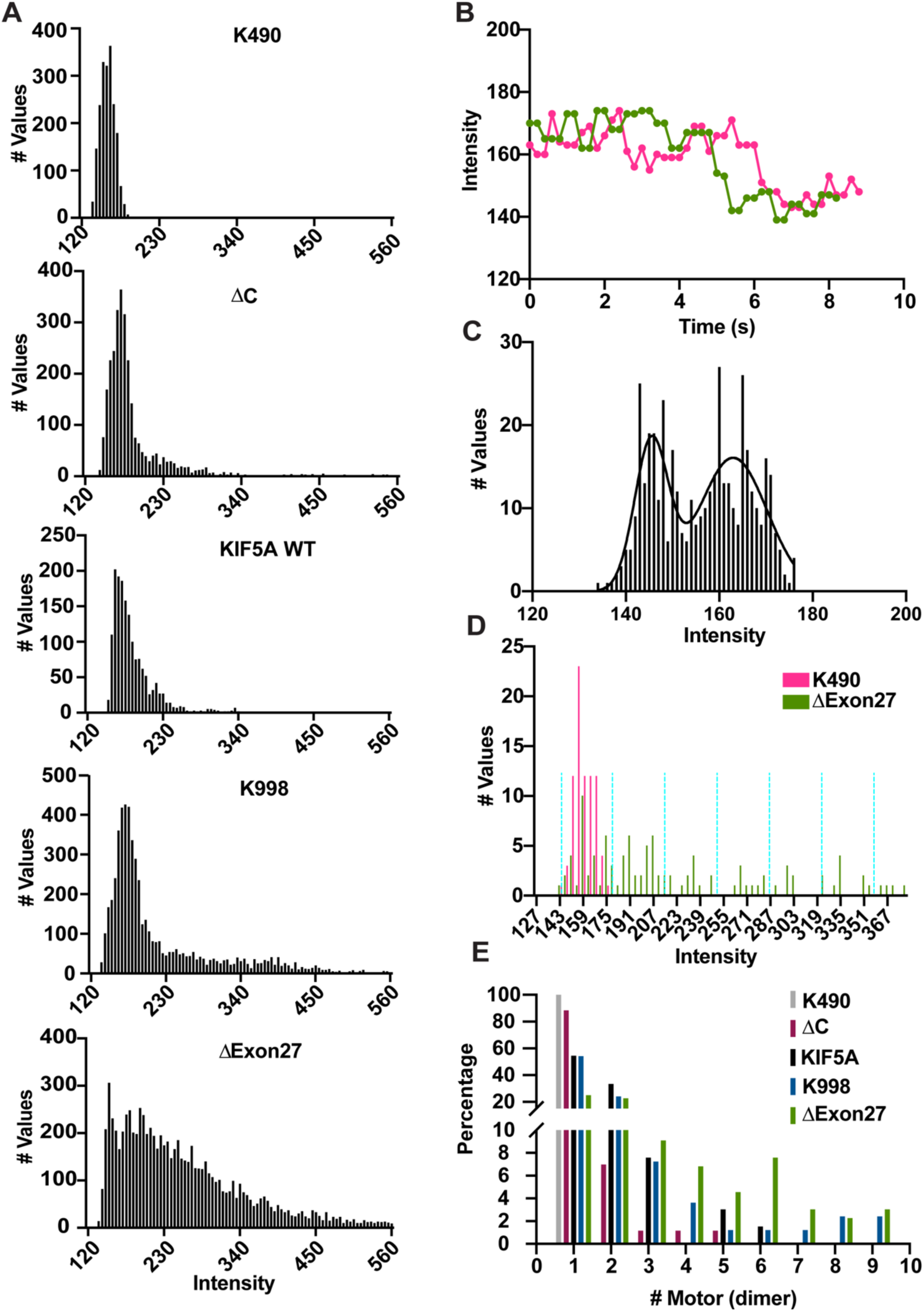
Enhanced motor self-association in KIF5A ΔExon27. **(A)** Histograms of the measured fluorescence intensities of the various moving kinesin constructs. The fluorescence intensity in every frame of each moving spot was measured, accumulated, and plotted in a histogram. Moving KIF5A ΔExon27 molecules exhibit a much larger fraction of high fluorescence intensities. The majority of the moving KIF5A ΔExon27 spots consist of multiple motors. (**B**) Two example traces of K490-EmGFP showing photobleaching events. K490-emGFP was used as the standard dimer molecule to determine the fluorescent intensity of EmGFP. (**C**) Histogram of measured intensities of moving K490-EmGFP molecules that showed a single photobleaching step. The intensity distribution was fit with two Gaussian functions, resulting in mean values of 145.5 ± 3.6 (mean ± SD) and 163.0 ± 7.3. With an average background fluorescence signal of 127.6 ± 0.7, one obtains an average fluorescent intensity is 17.7 for EmGFP (in a.u.). (**D-E**) Determination of the percentage of the number of motors for each motor construct. KIF5A K490 and KIF5A ΔExon27 are depicted here as examples **(D)**. The same method was applied to the other constructs. Each moving spot was assigned to a single intensity value that corresponded to the average intensity measured over the first 20 frames of the moving spot so the intensity would be less likely to be averaged down due to photobleaching. The measured values were then accumulated and plotted in a histogram. Based on the mean intensity of EmGFP, the measured intensity values were sorted into intervals of 35.4 (depicted by the turquoise dashed lines), with the first interval centered at 163 (163 ± 17.7). The percentage of each bin was calculated and then assigned to the calculated number of dimers (**E**). The percentages of a single dimer for all KIF5A constructs are: K490, 100%; ΔC, 88%; K998, 54%; WT, 54%; ΔExon27, 25%.

**Fig. S5.**
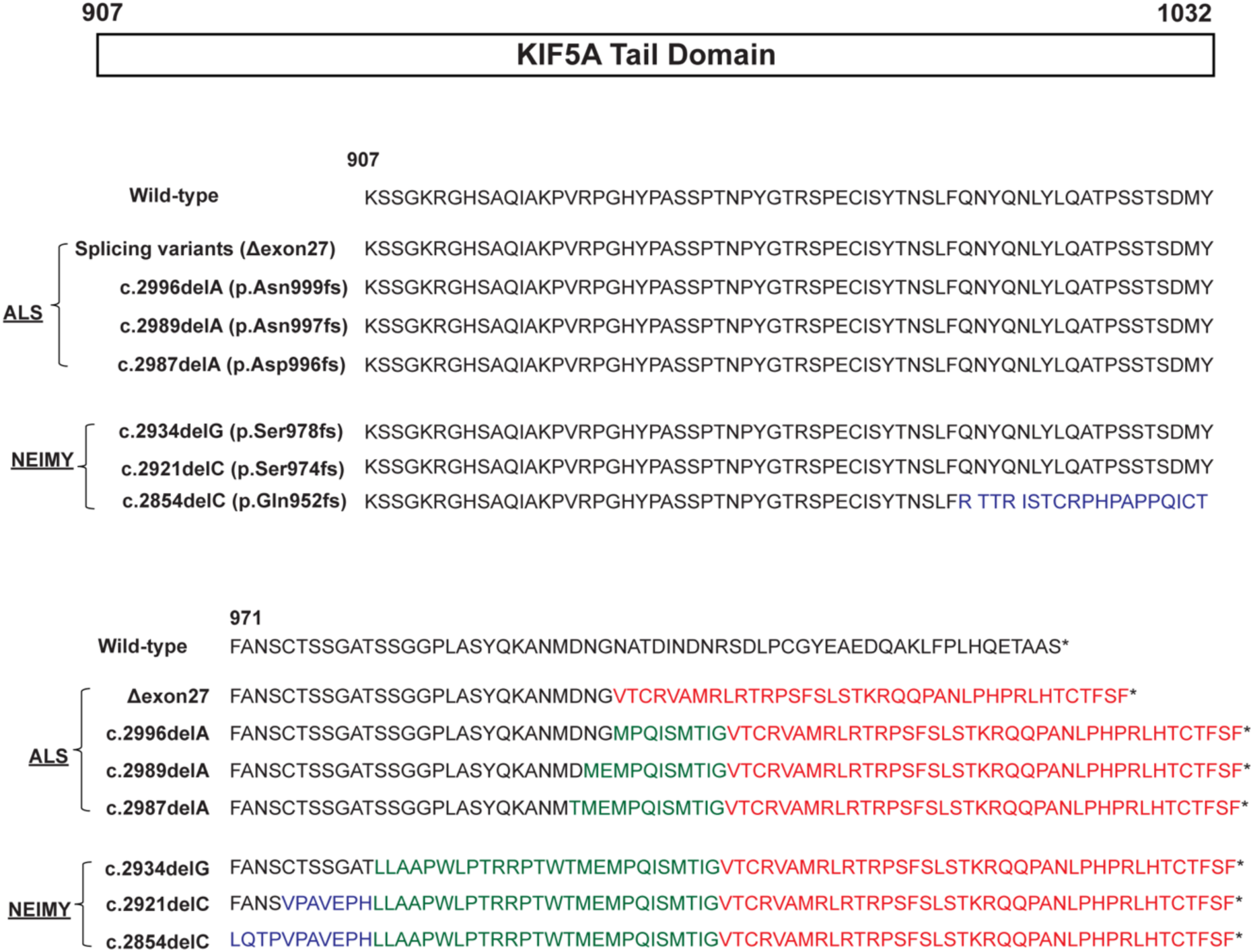
Comparison of amino acid sequences in the C-terminal tail domains of ALS-and NEIMY-associated KIF5A. The sequences in red are the new 39 amino acids of the C-terminal tail in ALS-associated KIF5A caused by mis-splicing of exon 27 (ΔExon27). The amino acids in blue and green are those different from WT caused by a single nucleotide deletion in three NEIMY cases.

**Fig. S6.**
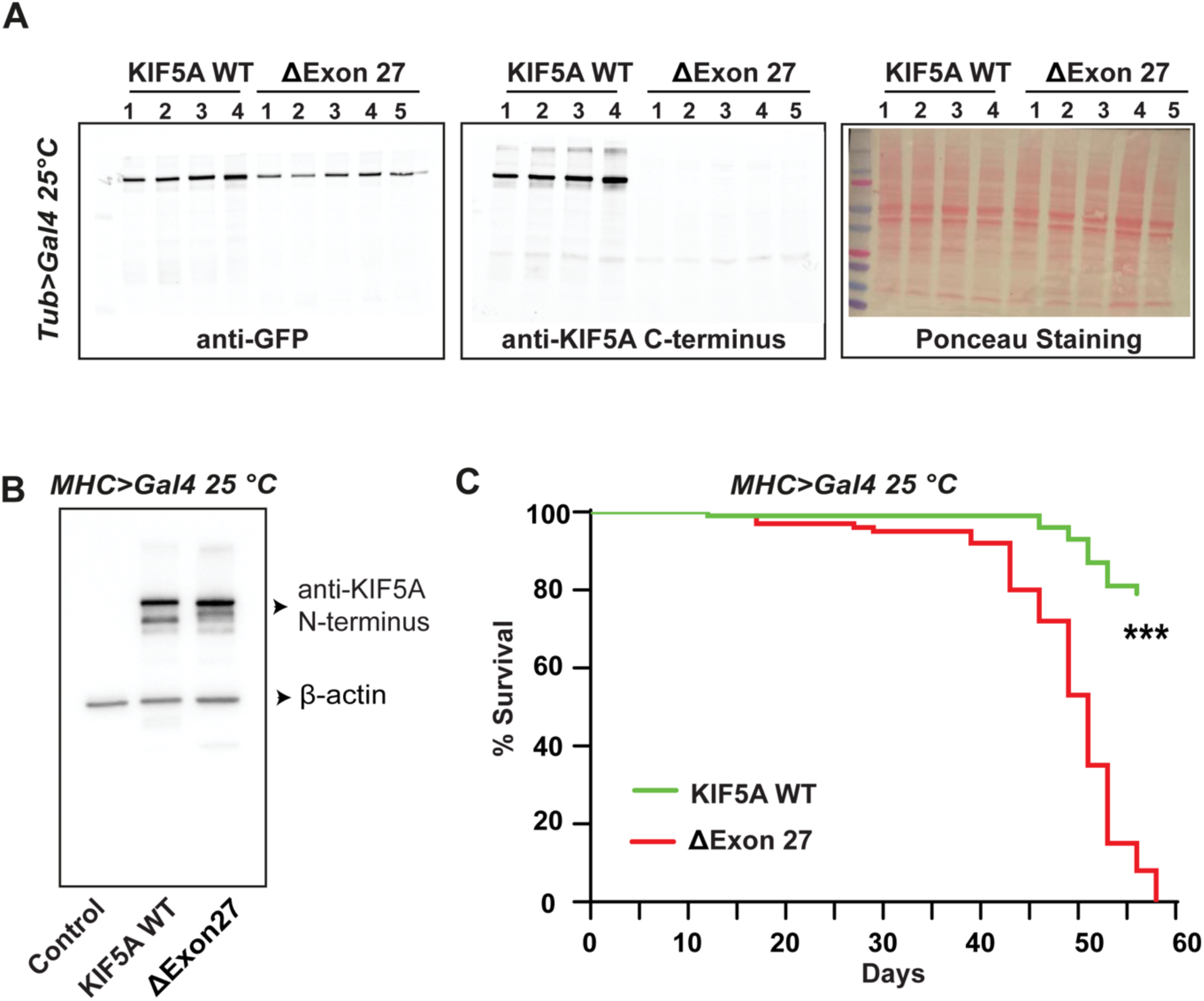
Muscle-specific expression of ΔExon27 in *Drosophila* leads to early lethality. **(A)** Western blot showing expression of human KIF5A proteins in *Drosophila* ubiquitously expressing WT or ΔExon27 driven by tubulin-Gal4. Genotypes were: KIF5A WT (UAS-KIF5A WT-GFP/ tubulin-Gal4); ΔExon27 (UAS-ΔExon27-GFP/ tubulin-Gal4). Samples from 4 (WT) and 5 (ΔExon27) different induvial lines were assessed with similar results. **(B)** Western blot showing expression of human KIF5A protein in *Drosophila* expressing WT or ΔExon27 in muscle tissues driven by MHC-Gal4. Genotypes were: Control (MHC-Gal4/+); KIF5A WT (UAS-KIF5A WT-GFP/MHC-Gal4); ΔExon27 (UAS-ΔExon27-GFP/MHC-Gal4) (**C**) Lifespan of male flies expressing KIF5A WT or ΔExon27 in muscles driven by MHC-Gal4 at 25 °C.

